# Maternal care during early development is necessary for the acquisition of a calming response to back stroking

**DOI:** 10.1101/2025.08.26.672261

**Authors:** Sachine Yoshida, Akiko Harauma, Toru Moriguchi, Yousuke Tsuneoka, Kimiya Narikiyo, Kazuya Miyanishi, Makoto Kashima, Makoto Wada, Yu Hayashi, Hiromasa Funato

## Abstract

In many mammals, early postnatal interactions between caregivers and offspring involve rich physical contact, including stroking, holding, and grooming. Offspring typically remain calm and close to the caregiver during such stimulation. Although these behaviors are thought to support emotional regulation and bonding during infancy, the underlying mechanisms and the role of prior experience remain unclear. Here, we demonstrate that back stroking induces a conserved calming response in both human infants and mouse pups, characterized by reduced movement and heart rate. In mouse pups, stroking further facilitated sleep onset, increased EEG delta power, and attenuated stress-induced corticosterone elevations. These sleep-promoting and stress-buffering effects were absent in artificially reared pups deprived of postnatal maternal contact, underscoring the importance of early tactile experience. Transcriptomic analysis revealed downregulation of the voltage-gated calcium channel subunit gene *Cacna1b* in the hypothalamus of artificially reared pups. Moreover, knockdown of hypothalamic *Cacna1b* in maternally reared pups abolished stroking-induced calming and sleep-promoting effects. This study identifies a conserved, experience-dependent calming response to affiliative tactile input, biologically embedded through plasticity, that supports physiological regulation and stress resilience during early development. Our results highlight that affiliative tactile sensation, like discriminative tactile sensation, depends on early experience to organize neural mechanisms regulating internal states.

**Significance:** Early-life physical contact with caregivers is essential for healthy emotional and physiological development in mammals, but its underlying mechanisms remain unclear. We show that back stroking induces a conserved calming response in both human infants and mouse pups, reducing movement, heart rate, and promoting sleep. These effects depend on early tactile experience and are mediated by *Cacna1b* expression in the hypothalamus. Our findings identify a plastic, experience-dependent pathway through which affiliative tactile input modulates internal states, revealing how parental care supports stress resilience and sleep physiology during early development.

## Introduction

Physical contact between caregivers and infants plays a critical role in supporting healthy emotional and physiological development. In humans, this principle is underscored by historical observations of infants raised in severely deprived conditions, such as institutionalized care with minimal caregiver interaction, who exhibited profound developmental delays and increased mortality rates (1, 2). These early findings have contributed to a widespread consensus that physical contact with caregivers is essential for optimal growth and well-being. The importance of early-life tactile input is not unique to humans. In many mammalian species, including rodents, maternal separation or early weaning leads to long-lasting behavioral and physiological changes in the offspring(3, 4). Despite the well-recognized benefits of caregiver–infant contact, the biological mechanisms through which affiliative tactile input supports emotional regulation and stress buffering remain incompletely understood. Although many studies have examined the adverse effects of tactile deprivation(1, 2, 5), fewer have focused on the potential adaptive functions of tactile input, such as its role in promoting relaxation and behavioral calmness.

Among the various forms of affiliative tactile input, stroking is a common and intentional behavior observed in daily caregiving practices. Parents often stroke their infant’s body to soothe distress or facilitate sleep(6). Similar behaviors have been documented in non-human mammals. For instance, macaque infants display fewer stress-related behaviors, such as scratching and agitation, after receiving stroking from caregivers(7). In rodents, maternal licking provides a form of tactile stimulation functionally analogous to stroking, and the amount of licking received in early life has been shown to influence stress responsiveness and emotional development in adulthood(8).

However, how maternal tactile input during early infancy shapes behavioral responses to somatosensory stimuli in later developmental stages has rarely been addressed and remain poorly understood. Furthermore, it remains unclear whether the calming effects of stroking are innate or shaped by postnatal tactile experience. The molecular and neural pathways that mediate these effects also require further clarification. To elucidate the developmental and molecular basis of this phenomenon, we examined the physiological and transcriptomic consequences of back stroking in both human infants and mouse pups, focusing specifically on the role of early-life tactile experience. We observed that back stroking reduced voluntary movement and heart rate in both species. In mice, this stimulation further promoted sleep onset and suppressed stress-induced hormone elevations. To evaluate the contribution of early tactile input, we compared artificially reared pups, which lacked maternal physical contact, with normally reared counterparts. We also conducted transcriptomic screening of the hypothalamus to identify genes influenced by early tactile experience and performed targeted knockdown experiments to assess their role in mediating the calming effects of stroking.

## Results

### Back stroking reduces voluntary movements and heart rate in human infants

To examine the real-time effects of mother’s stroking on their infants, we first investigated how maternal stroking affects the voluntary movement and heart rate of their infants. Each infant was attached to the disposable adhesive electrocardiogram (ECG) electrodes on their chests, and mother-infant pairs were video-recorded during the experiment sessions. Mothers were seated on a chair with their infant on their lap and instructed to gently but firmly stroke one of three body areas: the infant’s back, the back of the head, or the lower abdomen, for one minute each, in a randomized order. To minimize other verbal or nonverbal communication that can affect the infant’s responses, mothers were asked to refrain from making eye contact, speaking to their infant, or gently rocking them. Instead, mothers were instructed to stroke their infants with similar pressure and speed to what they usually do when soothing them. The sitting posture— whether the mother and infant faced each other or both faced forward—was left to the participants’ discretion. For mother–infant pairs who initially adopted a face-to-face position, the experiment began with stroking either the back or back-of-head (Fig. 1A). All analyses included only data from infants who did not cry during the procedures (7 males and 8 females for the back and lower abdomen stroking tasks; 5 males and 8 females for the back-of-head stroking task). Voluntary movements during the 1-minute stroking period were compared with those during the 1-minute pre-stroking period for each body part. Movements were categorized into three body regions: the head, upper body, and lower body. During the back-of-head stroking task, no significant changes in voluntary movement were observed in any body region compared to the pre-stroking period (Fig. 1B, left, Paired t-test with Holm correction, head: t = -1.53, df = 12, p = 0.15, upper body: t = 0.16, df = 12, p = 0.88, lower body: t = -1.72, df = 12, p = 0.11). Similarly, stroking of the lower abdomen did not result in significant changes in movement (Fig. 1B, middle, Paired t-test with Holm correction, head: t = 0.86, df = 14, p = 0.41, upper body: t = 0.94, df = 14, p = 0.36, lower body: t = 0.36, df = 14, p = 0.72). In contrast, during back stroking, voluntary movements in all three body regions were significantly reduced compared to the pre-stroking period (Fig. 1B, right, Paired t-test with Holm correction, head: t = 5.19, df = 14, p = 0.00014, upper body: t = 2.22, df = 14, p = 0.043, lower body: t = 3.88, df = 14, p = 0.0017). No significant sex differences were found in movement change scores (before – during stroking) across all three body areas (Welch’s t-test, p > 0.05, Table S1). Next, we compared the average heart rate before and during each stroking task. During back-of-head stroking, the heart rate significantly increased compared to the pre-stroking period (Fig. 1C, left, Paired t-test with Holm correction, t = -2.19, df = 12, p = 0.049). Although excluded from the analysis, the only infant who cried during the mother–infant sessions did so during back-of-head stroking. No significant difference in heart rate was observed during abdominal stroking (Fig. 1C, middle, Paired t-test with Holm correction, t = -1.76, df = 14, p = 0.10). In contrast, back stroking led to a significant decrease in heart rate relative to the pre-stroking period (Fig. 1C, right, Paired t-test with Holm correction, t = -2.24, df = 14, p = 0.042). There was no significant sex difference in heart rate across all three body areas (Welch’s t-test, p > 0.05, Table S1). These findings indicate the effects of maternal stroking vary depending on the body regions to be stroked. While stroking the back has a soothing effect on human infants, stroking the back of the head may have the opposite effect.

**Figure 1.**
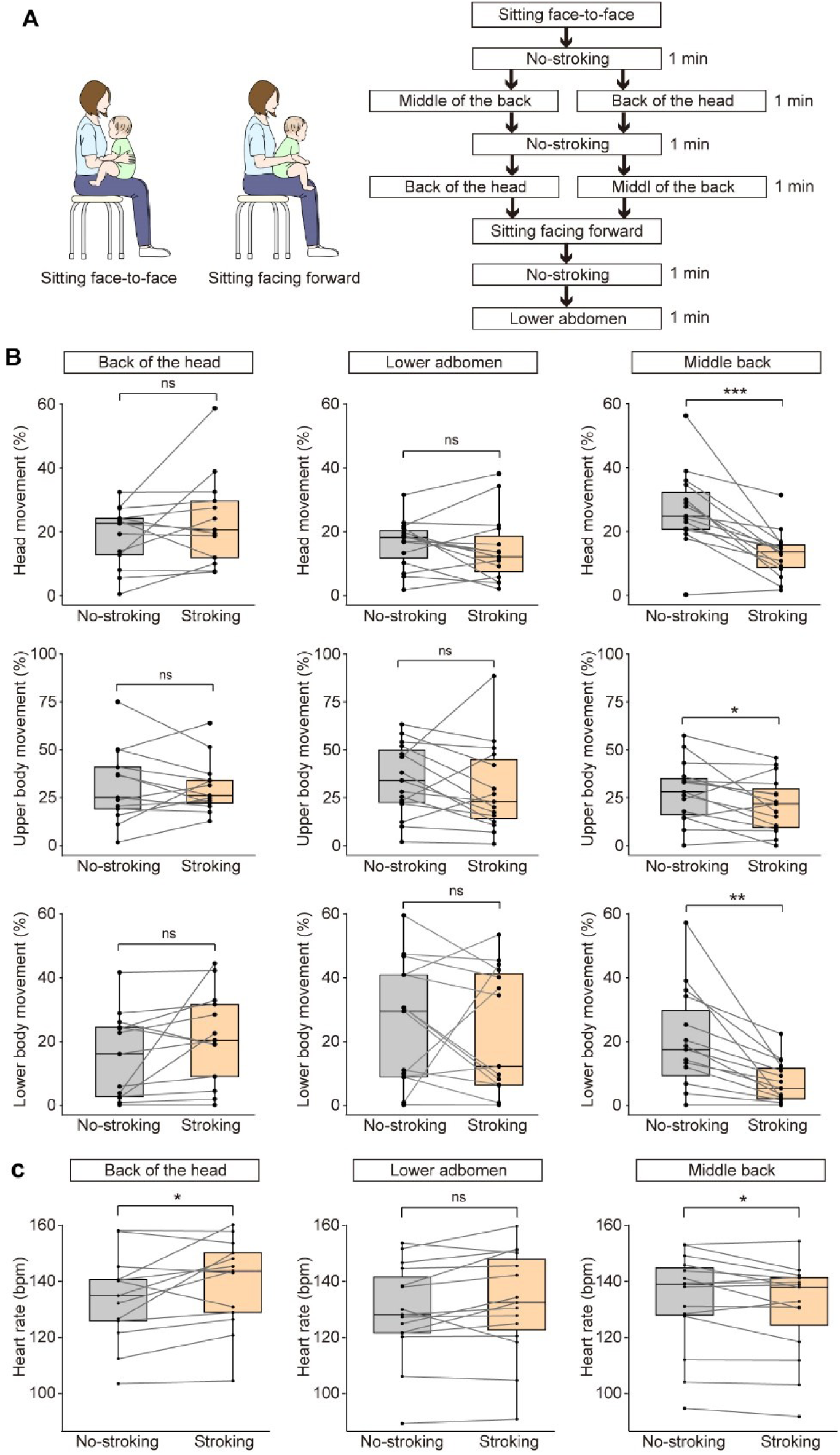
Calming responses to back stroking in human infants. (A) Illustration of two sitting positions used by mother–infant pairs (left) and experimental procedure when seated in a face-to-face position (right). (B) Comparison of spontaneous movements in the head, upper body, and lower body before and during stroking. The left panel shows data for the back of head stroking, middle for abdominal stroking, and right for back stroking. (C) Comparison of heart rate before and during stroking. The Left panel shows data for the back of head stroking, middle for abdominal stroking, and right for back stroking. The boxes represent the 25th, median, and 75th percentiles, and the whiskers represent the lowest or highest data within 1.53 interquartile range from the 25th or 75th percentile. n = 15 (m:7, f:8) in the lower abdomen and middle back stroking, and 13 (m:5, f:8) in the back of the head stroking. *: p < 0.05, **: p < 0.01. ns: not significant.

### Mouse dams lick the back of their pups more often than their chests

It is well known that the parents of many quadrupedal mammals, including rats and mice, frequently engage in licking behavior toward their offspring. This parental licking delivers tactile stimulation to the pups’ skin that resembles the sensory input induced by stroking. To investigate whether the reductions in spontaneous movement and heart rate observed in human infants in response to back stroking also occur in mouse pups, we first examined the preference of mouse dams for licking either the dorsal or ventral surfaces of their pups. Postnatal day (PND) 2 pups were removed from the home cage and marked on both the dorsal and ventral sides using a water-sensitive black ink that is removable through maternal licking. The brightness of each marked area was quantified both immediately before and four hours after the pups were returned to the home cage with their dams (Fig. 2A). The results showed that brightness significantly increased on the upper back, middle back, lower back, and chest, indicating that these areas had been licked (Fig. 2B, Paired t-test with Holm correction, upper back: t = -9.67, df = 8, p = 1.088e-05, middle back: t = -7.045, df = 8, p = 0.00011, lower back: t = -8.69, df = 8, p = 2.41e-05, chest: t = -4.60, df = 8, p = 0.0018). When comparing dorsal and ventral areas, all three dorsal areas exhibited significantly greater increases in brightness, suggesting that dams licked the dorsal side more frequently with their moist tongues (Fig. 2C, One-way ANOVA, F[3, 17.23] = 8.47, p = 0.001, Pairwise comparison with Holm correction, p = 0.0046 in upper back vs. chest, p = 0.020 in middle back vs. chest, p = 0.037 in lower back vs. chest).

**Figure 2.**
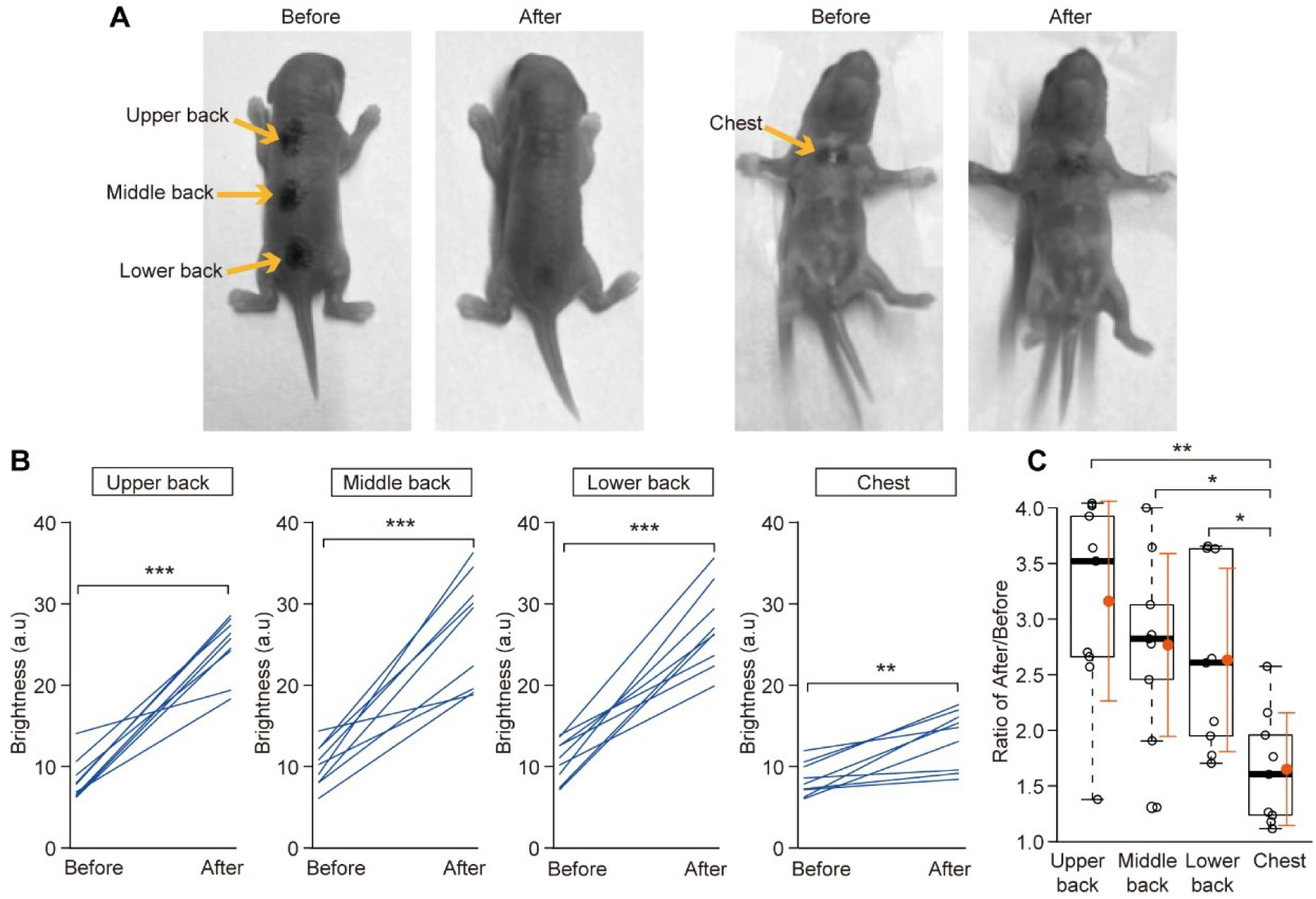
Region-specific quantification of maternal licking in neonatal mice. (A) Dorsal (left) and ventral (right) views of PND 2 mouse pups marked with ink. “Before” refers to the time point immediately after marking, prior to returning the pup to the home cage. “After” refers to four hours after the pup was returned to the home cage. (B) Within-individual changes in brightness for the upper back, middle back, lower back, and chest regions between before and after the 4-hour home cage exposure. (C) Comparison of brightness changes across different body regions. The boxes represent the 25th, median, and 75th percentiles, and the whiskers represent the lowest or highest data within 1.53 interquartile range from the 25th or 75th percentile. n = 4 (m:2, f:2). *: p < 0.05, **: p < 0.01, ***: p < 0.001.

### Back stroking mimicking the maternal licking induces NREM sleep in mouse pups

As the findings demonstrated that mouse pups frequently receive maternal contact on the back, we next examined the effect of back stroking on behavioral and physiological aspects in mouse pups. To isolate the influence of back stroking, pups were temporarily separated from their dam and gently restrained between soft packing foams. This setup allows for controlled stimulation of the back with a soft brush that mimics the dam’s licking behavior, while simultaneously recording electroencephalogram (EEG), ECG and electromyography (EMG) signals (Fig. 3A). At PND14, C57BL/6 (B6) pups made little attempt to escape and instead exhibited intermittent movements, such as pressing their noses into the corners or shifting their body orientation (Movie S1). A high-frequency band of EEG, such as 120–200 Hz or 110–300 Hz, contains components comparable to those of EMG, allowing for sleep-wake state identification(9, 10). Pups implanted with EEG electrodes and ECG wires were placed in a narrow enclosure, and underwent a 3-minute no-stroking period followed by a 3-minute back-stroking period (Fig. 3B). To elevate the pups’ arousal levels before each period, an unpleasant tactile stimulus was applied by pinna stroking for 1 minute. As a result, continuous back stroking led to a decrease in EMG activity and an increase in EEG delta power, resembling muscle tone and EEG patterns typically observed during non-rapid eye movement (NREM) sleep (Fig. 3B). Quantitative analysis revealed that continuous back stroking significantly decreased EMG activity and heart rate, while significantly increased delta power compared to the preceding no-stroking period (Fig. 3C, Paired t-test with Holm correction, EMG: t = 4.26, df = 5, p = 0.0080, heart rate: t = 3.52, df = 5, p = 0.017, delta power: t = -3.46, df = 5, p = 0.018). There was no significant sex difference in EMG, heart rate and delta power (Welch’s t-test, p > 0.05, Table S2). Furthermore, comparisons of delta power across periods revealed that delta power during stroking was significantly higher than during no-stroking period and reached levels comparable to those observed during NREM sleep in a home cage (Fig. 3D, A repeated-measures ANOVA, F [2, 12] = 43.03, p =3.36e-06, η²g = 0.511, Pairwise comparison with Holm correction, Stroking vs. No-stroking, p = 0.0030, NREM vs. No-stroking, p = 0.000046, Stroking vs. NREM, p = 0.056). No significant sex differences were found in the sum of root mean square (RMS) of EEG delta power (Welch’s t-test, p > 0.05, Table S3). Thus, continuous back stroking in mouse pups, like in human infants, suppressed voluntary movement, and decreased heart rate. Additionally, EEG recordings indicated that back stroking effectively induced NREM sleep in mouse pups.

**Figure 3.**
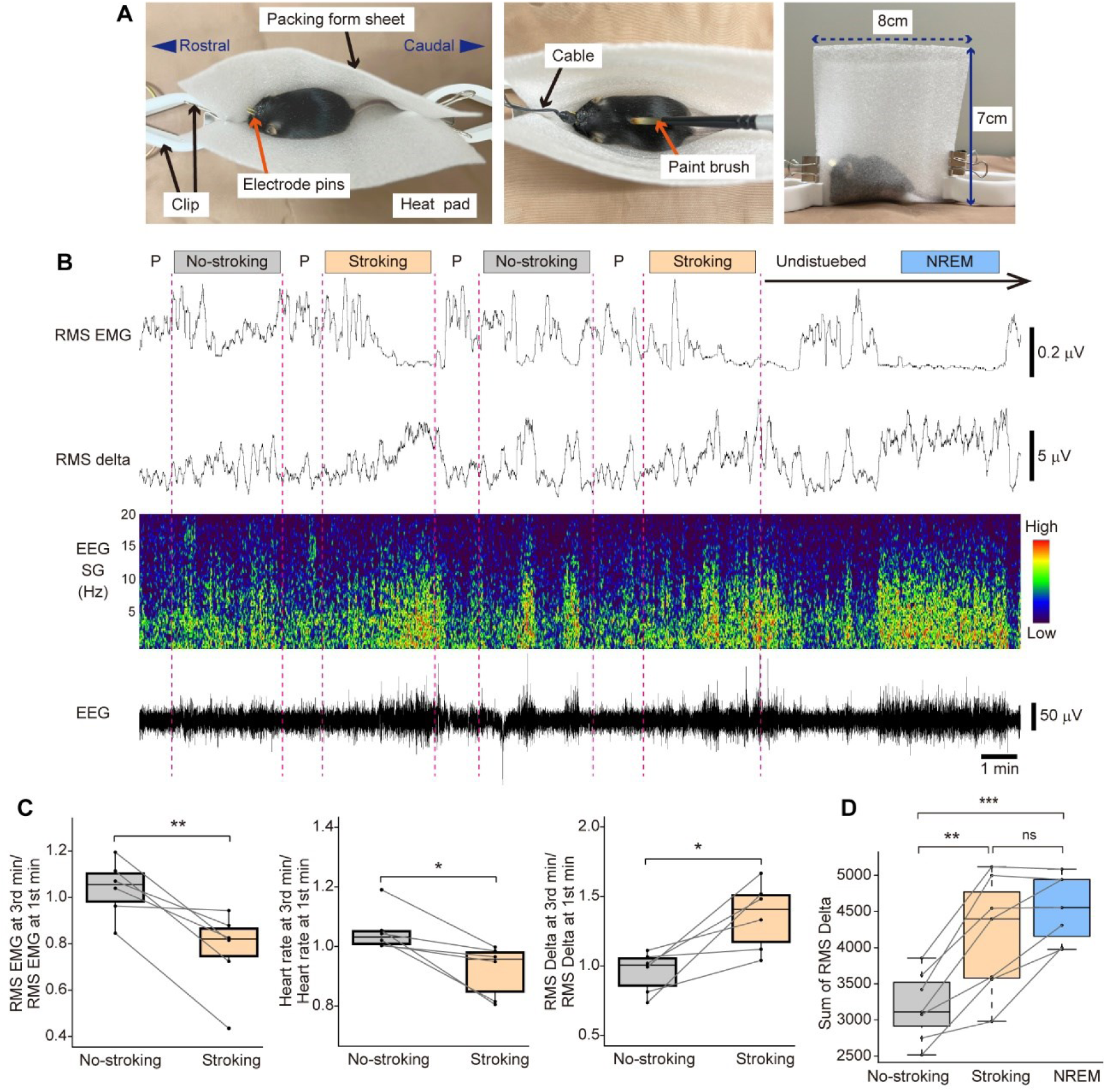
Calming responses to back stroking in mouse pups. (A) Three photographs of the experimental setup showing a gently restrained mouse pup undergoing simultaneous EEG, EMG, and ECG recordings during back stroking. (B) Representative traces showing changes across four conditions: pinna stroking (P), no-stroking period (No-stroking), back-stroking period (Stroking), and undisturbed period. Panels display (from top to bottom): RMS EMG, RMS delta power, EEG spectrogram (SG), and raw EEG waveform. (C) Comparison of the first and last 1-minute segments within 3-minute no-stroking and stroking periods. Plots show ratio changes in RMS EMG, heart rate, and RMS delta power. (D) Comparison of total RMS delta power during no-stroking, stroking, and NREM sleep periods. The boxes represent the 25th, median, and 75th percentiles, and the whiskers represent the lowest or highest data within 1.53 interquartile range from the 25th or 75th percentile. n = 6 (m:3, f:3) in (C) and n = 7 (m:4,f:3) in (D), *: p < 0.05, **: p < 0.01, ***: p < 0.001. ns: not significant.

### Back stroking has a stress buffering effect on mouse pups

Affiliative physical contact between parent and offspring plays a crucial role in the development of stress resilience(11). To examine whether back stroking confers stress-buffering effects, we conducted experiments using mouse pups. On PND14, pups were randomly assigned to one of three groups: (1) an isolation group, in which pups were placed alone in a narrow enclosure for 30 minutes (isolation stress); (2) an isolation-plus-stroking group, in which pups were placed alone in the enclosure and continuously stroked along the back with a brush throughout the 30-minute isolation period; and (3) an undisturbed control group, in which pups remained in the home cage with their dams and littermates without any experimental manipulation. Stress exposure is known to activate the hypothalamic–pituitary–adrenal (HPA) axis, eliciting endocrine responses via the activation of corticotropin-releasing hormone (CRH) neurons in the paraventricular nucleus (PVN) of the hypothalamus(12). The isolation group showed substantially higher expression of *c-Fos* mRNA, a marker of neuronal activity, in the PVN compared with the isolation-plus-stroking group (Fig. 4A). Consistently, the isolation group exhibited significantly higher corticosterone (CORT) levels compared to the other two groups at the end of 30 min session (Fig. 4B; One-way ANOVA, F[2, 5.59] = 46.39, p = 0.00033, Pairwise comparison with Holm correction, p = 0.00042 in isolation vs. isolation-plus-stroking, p = 0.0011 in isolation vs. undisturbed, p = 0.54 in undisturbed vs. isolation-plus-stroking). In contrast, CORT levels in the isolation-plus-stroking group were comparable to those in the undisturbed group, with no significant difference (Fig. 4B). No significant sex differences were found in each group (2-3 males, 2 females, Welch’s t-test, p > 0.05, Table S4). Similar results were observed in pups at PND11, which corresponds to the late phase of the stress hyporesponsive period(13), indicating that the stress-buffering effect of stroking emerges prior to the end of this period (Figure S2). As for oxytocin, a hormone involving in the formation of social bonds(14), no significant differences in its concentration were observed among the three groups on PND14. (Fig.4C; One-way ANOVA, F[2, 3.75] = 0.033, p = 0.97). Together, these findings demonstrate that affiliative tactile stimulation induced by back stroking has a stress-buffering effect in developing mice.

**Figure 4.**
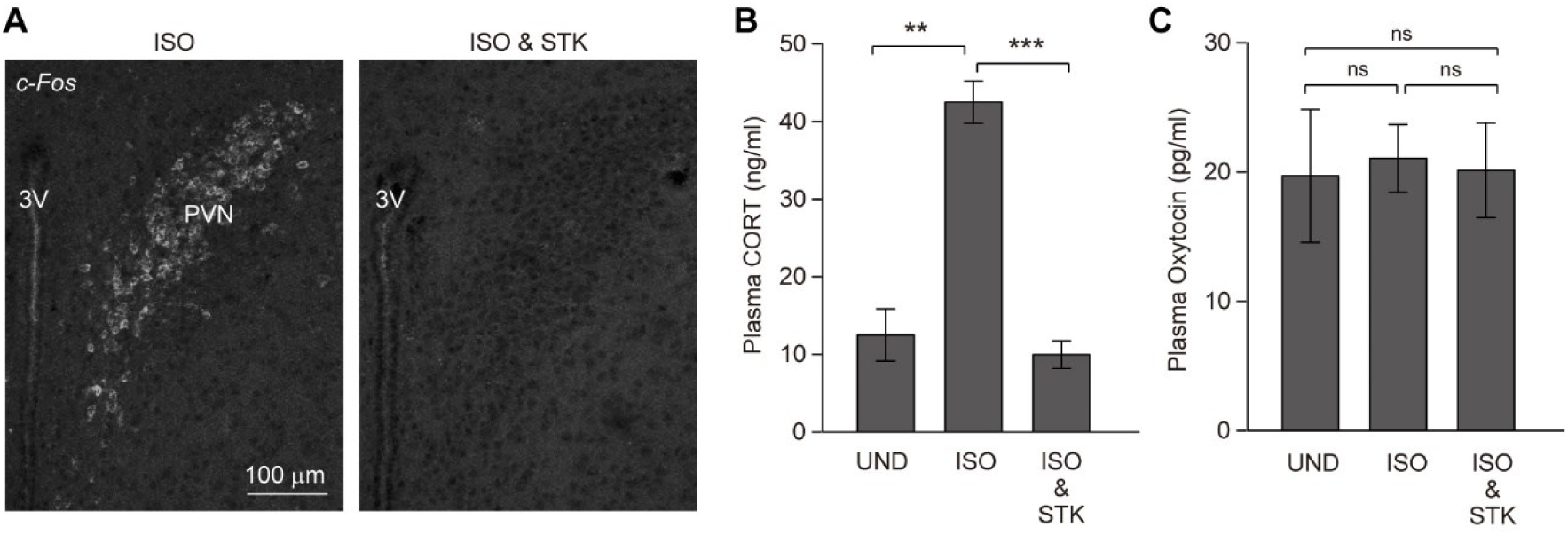
Back stroking acts as a stress-buffering stimulus in mouse pups. (A) Comparison of *c-Fos* mRNA expression in the paraventricular nucleus (PVN) of the hypothalamus between isolated pups (ISO) and pups stroked during isolation (ISO & STK). (B) Plasma corticosterone (CORT) levels in pups under undisturbed conditions (UD), isolation, or stroking during isolation. (C) Plasma oxytocin levels in pups under undisturbed conditions, isolation, or stroking during isolation. UD: n = 5 (m: 3, f: 2), ISO: n = 4 (m: 2, f: 2), STK: n = 4 (m: 2, f: 2). 3V, third ventricle, PVN: paraventricular nucleus. Error bars represent mean ± SD. **: p < 0.01, ***: p < 0.001. ns: not significant.

### Artificially reared pups fail to show drowsiness and stress buffering effect induced by the back stroking

During infancy, mammals exhibit various innate reflexes, such as the sucking reflex, in which pups begin to suck in response to tactile stimulation on the roof of the mouth. Fixed motor or postural responses to specific sensory stimuli are not limited to the very early stage of development. For example, Transport Response emerges from PND8 to PND16(15). These findings prompted us to examine whether the calming effect of back stroking is innate or acquired through early-life tactile experiences, particularly those provided by maternal caregiving. To deprive pups of maternal licking and other parental gentle contact, we employed artificially reared (AR) protocol with scheduled milk-feeding during the first two weeks after birth. We then compared their responses to back stroking with those of maternally reared (MR) pups by assessing changes in EMG, heart rate, EEG delta power and plasma CORT levels.

During the optimization of AR conditions, we initially used C57BL/6 mice or C57BL/6 × ICR hybrids. However, pups from these strains consistently failed to survive beyond the first postnatal week when reared artificially, indicating that the artificial rearing is particularly challenging in these backgrounds. We therefore selected the inbred ICR strain for all subsequent experiments due to its high survival rate, robust milk consumption, and consistent weight gain under artificial rearing conditions (Fig. 5A, Movie S2). Each feeding session included genital stimulation to induce urination and defecation, followed by weighing before and after milk ingestion. Milk intake was deemed complete when the pup spontaneously rejected the custom-designed artificial nipple. Throughout the rearing period, experimenters wore gloves, minimized handling, and refrained from grooming or stroking the pups. Our artificial rearing protocol supported overall weight gain comparable to that of the MR group (Fig. 5B; Welch’s t-test: PND2, t = 2.63, df = 12.70, p = 0.021; PND4, t = 1.19, df = 14.22, p = 0.26; PND6, t = –3.78, df = 11.97, p = 0.0026; PND8, t = 0.72, df = 18.68, p = 0.48; PND10, t = –1.37, df = 18.96, p = 0.19; PND12, t = –1.85, df = 23.39, p = 0.077). AR pups, which are not licked by their dams, often have fur soiled with milk and feces and appear unkempt, but they exhibit no apparent motor abnormalities. Eyelid opening in AR pups occurred from PND12 to PND13, which was comparable to that in MR pups. There were no significant differences between MR and AR pups in the latency to perform the righting reflex at PND12 or in the duration of immobilization during the Transport Response at PND13, a filial-specific calming reaction observed during parent–infant transport in mammals(16) (Fig. S3A and B, righting reflex: Welch’s t-test: t = 0.38, df = 11.94, p = 0.71, Transport Response: Welch’s t-test: t = 0.15, df = 9.99, p = 0.88). A small number of pups were raised beyond weaning to assess potential behavioral abnormalities. At 8 weeks of age, no significant differences were observed between MR and AR mice in immobility time during a 6-minute tail suspension test, a widely used behavioral assay for depressive-like behavior (17) (Fig. S3C, Welch’s t-test, t = –0.33, df = 5.59, p = 0.75, n = 4 per group).

**Figure 5.**
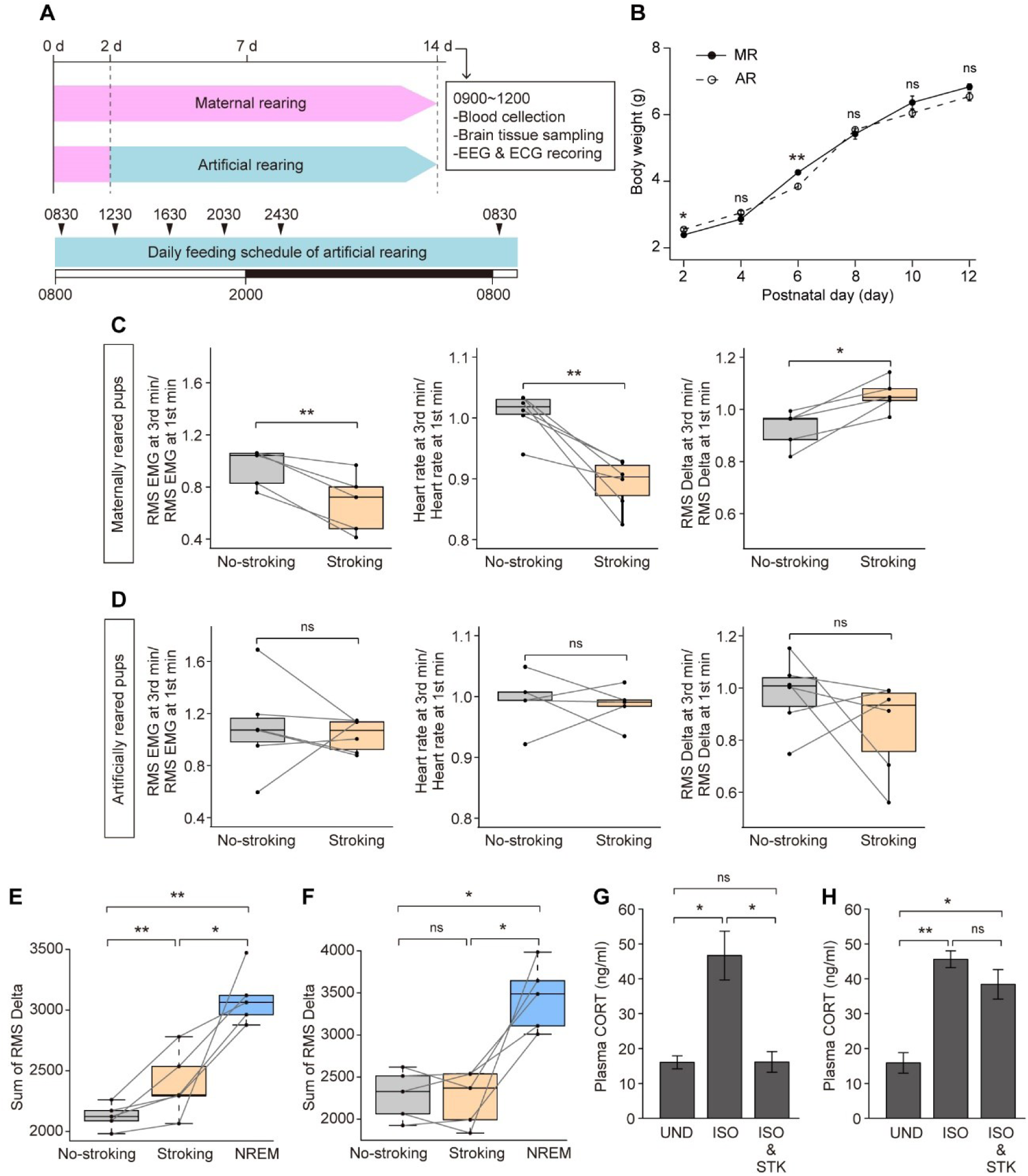
Stroking fails to induce sleep onset or stress buffering in artificially reared pups. (A) Experimental timeline for artificially reared pups, including scheduled feeding times and the sequence of behavioral and physiological assessments. White and black bars indicate the light and dark phases, respectively. (B) Comparison of body weight between maternally reared (MR) and artificially reared (AR) pups. (C, D) Comparison of the first and last 1-minute segments within 3-minute no-stroking and stroking periods. Plots show ratio changes in RMS EMG, heart rate, and RMS delta power in MR (C) and AR pups (D). In (C), RMS EMG, heart rate, and RMS delta power, n = 5 (3 males, 2 females). In (D), EMG and delta power, n = 6 (m: 3, f: 3), except heart rate, n = 4 (m: 2, f: 2) due to ECG wire disconnection. (E, F) Comparison of total RMS delta power during no-stroking, stroking, and NREM sleep periods in MR (E) and AR (F) pups. n = 5 (m: 3, f: 2) and n = 6 (m: 3, f: 3) in (E) and (F), respectively. (G, H) Plasma CORT levels in MR (G) and AR (H) pups under undisturbed conditions (UD), isolation (ISO), or stroking during isolation (ISO & STK). The boxes represent the 25th, median, and 75th percentiles, and the whiskers represent the lowest or highest data within 1.53 interquartile range from the 25th or 75th percentile. n = 4–5 (m: 2–3, f: 2) per group and n = 3–4 (m: 1–2, f: 2) per group in (G) and (H), respectively. Error bars represent mean ± SD. *: p < 0.05, **: p < 0.01. ns: not significant.

On the morning of PND 13, we placed both AR and MR ICR pups in a narrow enclosure as performed with C57BL/6 pups and recorded EMG, heart rate and EEG during no-stroking and back stroking periods. In MR pups, as observed in maternally reared C57BL/6 pups, back stroking significantly reduced EMG and heart rate and increased delta power compared to the preceding no-stroking periods (Fig. 5C, Paired t-test with Holm correction, EMG: t = 5.079, df = 4, p = 0.0071, heart rate: t = 4.72, df = 5, p = 0.0052, delta power: t = -4.55, df = 4, p = 0.010). In contrast, AR pups showed no such reduction in EMG or heart rate, nor any increase in delta power upon back stroking (Fig. 5D, Paired t-test with Holm correction, EMG: t = 0.43, df = 5, p = 0.68, heart rate: t = 0.30, df = 4, p = 0.78, delta power: t = 1.13, df = 5, p = 0.31). Although the delta power during stroking was smaller than that observed during NREM sleep in MR pups, it was significantly higher than during the no-stroking condition, indicating that stroking stimulation induces a drowsy state (Fig. 5E, A repeated-measures ANOVA, F [2, 8] = 24.27, p = 4.010e-04, η²g = 0.82, Pairwise comparison with Holm correction, Stroking vs. No-stroking, p = 0.040, NREM vs. No-stroking, p = 0.0070, Stroking vs. NREM, p = 0.040). In AR pups, delta power during both no-stroking and stroking periods was similar, and both were significantly lower than delta power during NREM sleep (Fig. 5F, A repeated-measures ANOVA, F [2, 8] = 22.25, p = 5.39e-04, η²g = 0.77, Pairwise comparison with Holm correction, Stroking vs. No-stroking, p = 0.712, NREM vs. No-stroking, p = 0.017, Stroking vs. NREM, p = 0.022). In both AR and MR pups, no significant sex differences were found in EMG, heart rate, delta power, or the sum of RMS delta (Welch’s t-test, p > 0.05, Table S2 and S3).

We next examined whether 30 minutes of continuous back stroking could buffer isolation-induced stress in MR and AR pups. As observed in C57BL/6 pups, maternally reared ICR pups subjected to stroking during isolation showed CORT levels comparable to those of undisturbed controls, indicating effective stress buffering (Fig. 5G; One-way ANOVA, F [2, 5.28] = 8.049, p = 0.025, Pairwise comparison with Holm correction, p = 0.0047 in isolation vs. isolation-plus-stroking, p = 0.0047 in isolation vs. undisturbed, p = 0.98 in undisturbed vs. isolation-plus-stroking). In contrast, AR pups showed no such reduction in CORT levels. Their CORT concentrations remained as elevated as those of AR pups subjected to isolation without stroking (Fig. 5H; One-way ANOVA, F [2, 5.28] = 27.69, p = 0.0016, Pairwise comparison with Holm correction, p = 0.20 in isolation vs. isolation-plus-stroking, p = 0.0016 in isolation vs. undisturbed, p = 0.012 in undisturbed vs. isolation-plus-stroking). In both AR and MR pups, no significant sex differences were found in each group (Welch’s t-test, p > 0.05, Table S4).

Taking together, these results demonstrate that early-life exposure to maternal licking and all other forms of tactile stimulation is essential for the development of physio-behavioral responses to stroking. In the absence of such early tactile experiences, back stroking fails to induce drowsiness or attenuate stress responses in developing mice.

### *Cacna1b* knockdown pups fail to show drowsiness induced by the back stroking

Next, we investigated differentially expressed genes between MR and AR pups in ICR strains. At PND13, total RNA was extracted from the hypothalamic region, given its pivotal role in sleep–wake regulation(18) and stress responses(19). Among the 55414 genes analyzed, 20 genes were differentially expressed (FDR = 0.05) (Fig. 6A, Fig.S4, Table S5). Among these, we focused on the *Cacna1b* gene, which encodes the Cav2.2 voltage-gated calcium channel subunit, due to its essential role in synaptic neurotransmitter release in the central nervous system(20) and its reported hyperactivation phenotype in knockout mice(21).

**Figure 6.**
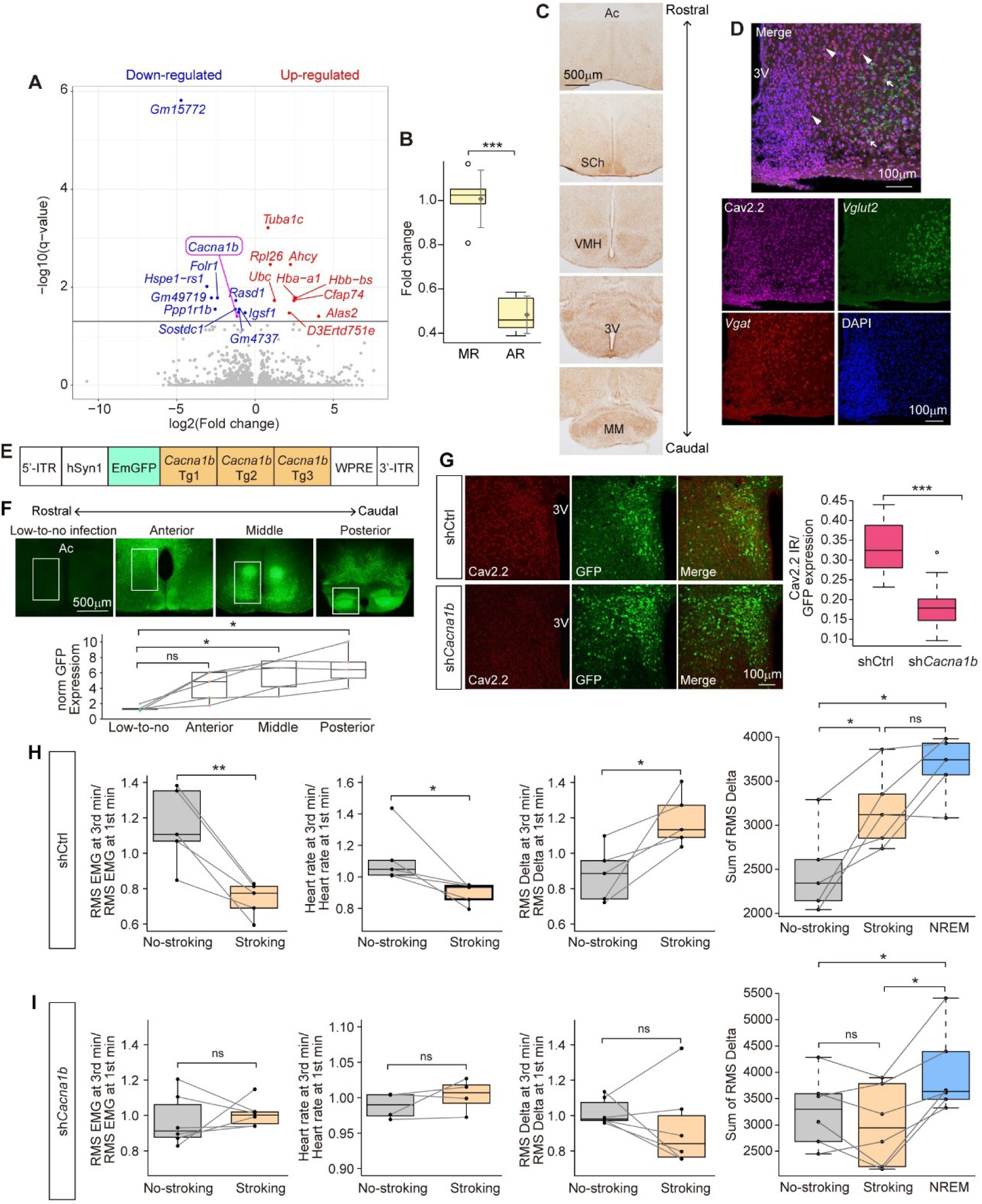
Hypothalamic *Cacna1b* expression regulates stroking-induced calming response in mouse pups. (A) Volcano plot showing differentially expressed genes in the hypothalamus of maternally reared (MR) and artificially reared (AR) pups based on RNA-seq analysis. n=4 and 3 in MR and AR, respectively. (B) Quantitative PCR comparison of hypothalamic *Cacna1b* expression between MR and AR pups. n = 5 in each group. (C) Immunohistochemistry for Cav2.2, encoded by the *Cacna1b* gene, in the hypothalamus. (D) Combined immunohistochemistry and isHCR detecting Cav2.2 protein, *Vglut2* mRNA, and *Vgat* mRNA in the hypothalamus. Arrows and arrowheads indicate Cav2.2/*Vglut2* and Cav2.2/*Vgat* double-positive neurons, respectively. (E) Schematic of the AAV vector carrying a construct for knockdown of the *Cacna1b* gene. (F) Expression and regional distribution of the *Cacna1b* knockdown vector in the hypothalamus visualized by GFP (top). Quantification of GFP expression in the non-infected area and in three infected hypothalamic regions (anterior, middle, and posterior parts) (bottom). Norm GFP indicates GFP expression normalized to fluorescence intensity measured in a non-infected cortical region. (G) GFP expression and Cav2.2 immunoreactivity (IR) in the hypothalamus of pups injected with shCtrl or sh*Cacna1b* vectors (left), and quantification of Cav2.2 immunoreactivity (right). n = 22 cells prepared from two pups of each rearing group. (H, I) Comparison of the first and last 1-minute segments within 3-minute no-stroking and stroking periods. Plots show ratio changes in RMS EMG, heart rate, and RMS delta power in pups with shCtrl (H) or sh*Cacna1b* (I). Sum of RMS delta power during no-stroking, stroking, and NREM sleep conditions in pups with shCtrl (H) or sh*Cacna1b* (I). In (H), RMS EMG, heart rate, RMS delta n = 5 (m: 2, f: 3). In (I), RMS EMG and RMS delta, n =6 (m: 3, f: 3), heart rate n = 4 (m: 2, f: 2). The boxes represent the 25th, median, and 75th percentiles, and the whiskers represent the lowest or highest data within 1.53 interquartile range from the 25th or 75th percentile. 3V: third ventricle, AC: anterior commissure, MM: mammillary nucleus, SCh: suprachiasmatic nucleus, VMH: ventromedial hypothalamic nucleus. *: p < 0.05, **: p < 0.01, ***: p < 0.001. ns: not significant.

Notably, *Cacna1b* mRNA exhibits a distinct temporal expression pattern during development, with elevated and widespread expression across the brain during the first two postnatal weeks, followed by a clear decline at the fourth week after weaning(22).

RNA-seq analysis revealed a 45.14% reduction in *Cacna1b* mRNA expression in AR pups compared to MR pups (Fig.6A, log_2_FC = -1.15, *q*-value = 3.93e^-2^). Quantitative RT-PCR using total RNA from ICR pups confirmed this result, showing an approximately 48% reduction in *Cacna1b* mRNA expression in AR pups (Fig. 6B; Welch’s t-test: t = 7.54, df = 6.89, p = 0.00014). Immunohistochemical analysis using PND14 C57BL/6 brain sections revealed broad Cav2.2 immunoreactivity throughout the hypothalamic region used for RNA extraction, spanning from the preoptic area to the retromammillary nucleus (Fig. 6C). Double staining demonstrated Cav2.2 expression in both glutamatergic and GABAergic neurons (Fig. 6D) as reported in previous studies(23).

To examine the functional consequence of *Cacna1b* gene downregulation, we generated a short hairpin RNA (shRNA) plasmid construct (sh*Cacna1b*) containing three distinct target sequences (Fig. 6E, Table S6). C57BL/6 pups were bilaterally injected with the viral vector into the hypothalamus at PND1 and examined at PND14. At PND14, the present stereotaxic coordinates yielded widespread GFP expression throughout hypothalamic areas (Fig. 6F). We quantitatively assessed the knockdown efficiency of *Cacna1b* by analyzing immunohistochemical signal intensity in pups injected with either non-silencing control shRNA (shCtrl) or sh*Cacna1b*. In sh*Cacna1b* pups, Cav2.2 immunoreactivity, normalized to GFP expression, was reduced to nearly half, indicating that sh*Cacna1b* effectively downregulated Cav2.2 protein expression (Fig. 6G; Welch’s t-test: t = 8.83, df = 40.32, p = 5.673e-11). We next examined whether back stroking in sh*Cacna1b* pups would induce reductions in voluntary movement and heart rate, as well as an increase in delta power. In shCtrl pups, as observed in maternally reared C57BL/6 and ICR pups, continuous back stroking significantly reduced EMG activity and heart rate, while significantly increasing delta power compared to the preceding no-stroking period (Fig. 6H; paired t-test with Holm correction: EMG: t = 5.19, df = 4, p = 0.0065; heart rate: t = 2.81, df = 4, p = 0.048; delta power: t = -3.23, df = 4, p = 0.032). Delta power during stroking was significantly higher than during the no-stroking period and reached levels comparable to those during NREM sleep (Fig. 6H; repeated-measures ANOVA: F [2, 8] = 18.94, p = 9.25e-04, η²g = 0.60; pairwise comparisons with Holm correction: stroking vs. no-stroking, p = 0.042; NREM vs. no-stroking, p = 0.11; stroking vs. NREM, p = 0.069).

In contrast, sh*Cacna1b* pups showed no reduction in EMG activity or heart rate, nor any increase in delta power following back stroking (Fig. 6I; paired t-test with Holm correction: EMG: t = -0.52, df = 5, p = 0.63; heart rate: t = -1.19, df = 3, p = 0.32; delta power: t = 0.93, df = 5, p = 0.39). Furthermore, in sh*Cacna1b* pups, delta power during both no-stroking and stroking periods remained comparable and significantly lower than that observed during NREM sleep (Fig. 6I; repeated-measures ANOVA: F [2, 10] = 16.40, p = 6.96e-04, η²g = 0.27; pairwise comparisons with Holm correction: stroking vs. no-stroking, p = 0.18; NREM vs. no-stroking, p = 0.013; stroking vs. NREM, p = 0.013). These results indicate that somatosensory stimulation from maternal licking during early postnatal development contributes to the maintenance of Cav2.2 channel expression in the hypothalamus, and that Cav2.2-mediated neural signaling plays a critical role in the expression of NREM sleep induced by gentle back stroking.

## Discussion

Here, we demonstrated that affiliative stroking stimulation rapidly induces calming responses in both human infants and mouse pups. This stimulation resulted in reduced spontaneous movements and lowered heart rates. In mice, it additionally promoted the onset of sleep and reduced stress levels (Figures 1, 3, 4). These effects were dependent on early-life physical contact with the dam. Mouse pups that were artificially reared without maternal contact did not show calming or sleep-promoting responses to stroking. Instead, they exhibited elevated levels of stress markers (Figure 5). At the molecular level, this developmental deficit was associated with reduced expression of *Cacna1b*, the gene that encodes the Cav2.2 voltage-gated calcium channel, in the hypothalamus. Furthermore, targeted knockdown of *Cacna1b* in the hypothalamus of maternally reared pups eliminated the calming and sleep-inducing effects of stroking (Figure 6). These results suggest the presence of a conserved calming mechanism induced by affiliative tactile stimulation in mammals.

Neurological reflexes and behavioral responses that are fixed motor or behavioral reactions to specific sensory stimuli temporarily appear during infancy. These responses are essential for pup survival and facilitate parental caregiving behavior. For example, the sucking reflex, in which pups begin to suck in response to tactile stimulation on the roof of the mouth, enables pups to obtain milk from their mother’s nipples. Similarly, the Transport Response that is a postural reaction elicited by pressure on the upper back, emerge between PND8 and PND16. This response helps parents safely carry their pups as the pups grow in size and weight(16). In a similar developmental period, calming responses to back stroking are observed, which reduces pup activity and likely eases maternal labor by temporarily pacifying the pups. As weaning approaches, pups begin to behave more independently and forage for food in the home cage without relying on their dams. At this stage, back stroking no longer induces a calming effect and may even provoke agitation. Adult mice do not exhibit calming responses to back stroking by experimenters without prior training(24), suggesting that this behavioral response is developmentally regulated.

Although developmentally regulated reflexes and behavioral responses are generally innate and occur rapidly in response to sensory stimuli, the calming response to back stroking is distinct in that it takes minutes to develop and is acquired through parental care. Unlike rapid reflex arcs that involve direct pathways from sensory inputs to motor neurons, the calming effect may require sensory information to ascend to higher brain regions, where it modulates neural networks regulating sleep/wake state, particularly within the hypothalamus and other brain areas. Our current findings support the notion that the development of somatosensory-related reflexes and responses, such as the righting reflex and the Transport Response, can proceed largely independent of direct maternal contact in the early postnatal period. Previous studies have shown that fundamental aspects of somatosensation processing are established prenatally and continue to mature after birth(25). Despite being deprived of maternal tactile interactions, artificially reared mice exhibited normal expression of the sucking reflex, righting reflex, and Transport Response. These mice also showed body weight growth comparable to those of maternal reared mice and no apparent abnormalities in sleep/wake behavior in the home cage. While we cannot rule out the possibility that artificial rearing disrupts several aspects of neural development, our data indicate that key sensory-motor responses are preserved under artificially rearing conditions.

Although gentle touch on the skin elicits pleasant sensations via unmyelinated C-fiber low threshold mechanoreceptors (C-LTMRs)(26), the present human and mouse studies involves stronger pressure than typical gentle touch. Human caregivers were instructed to stroke the infant’s back with firm pressure to ensure that the stimulation was effectively conveyed. In mouse pups, back stroking with a brush mimicked the pressure exerted by maternal licking and grooming. Adequate pressure is necessary for mothers to remove milk and excreta stains from the pup’s body. Through artificial rearing experiments, we confirmed that gentle touch or light brushing alone is not sufficient to remove such stains from the body surface of artificially reared pups. Therefore, maternally reared pups are frequently subjected to stroking that may activate mechanoreceptors not only in the dermis but also in the subcutaneous tissue. Such painless, composite tactile pressure information is conveyed to the brain via spinal projection neurons (SPNs) in the dorsal horn. SPNs send direct and indirect projections to multiple brain regions, including broad areas of the hypothalamus(27). The hypothalamus serves as a central integrative hub that regulates homeostasis and instinctive behaviors, including sleep/wakefulness and stress responses(18). It is therefore likely that stroking the back modulates neuronal activity in the PVN of the hypothalamus either directly or indirectly, which may trigger a systemic calming response that integrates neuroendocrine, autonomic, and sleep–wake regulation in the offspring. Consistently, back stroking led to reduced heart rate and lowered circulating CORT levels, suggesting a shift toward parasympathetic dominance and decreased HPA axis activity. However, back stroking did not alter plasma oxytocin levels, that implicated in neural responses to social bonding and social touch (28, 29). These physiological changes may, in turn, facilitate the induction of NREMS-like state.

Consistent with the role of the hypothalamus in regulating sleep/wakefulness and stress responses, which are associated with the calming effects of back stroking (18), artificially reared mice showed altered gene expression profiles in the hypothalamus at PND13. Artificially reared mice showed reduced expression of *Cacna1b*, which encodes the core subunit of the N-type voltage-gated calcium channel, Cav2.2, a channel known to mediate presynaptic neurotransmitter release. Furthermore, knockdown of *Cacna1b* in the hypothalamus in maternally reared mice abolished the calming response to back stroking. These findings suggest that maternal caregiving may help maintain sufficient level of *Cacna1b* expression in the hypothalamus, thereby facilitating the postnatal development of calming response to back stroking. However, since our genetic manipulation broadly affects the hypothalamus and *Cacna1b* is widely expressed throughout the hypothalamus, the specific neural populations or circuitries mediating this effect remain unidentified. Although the functional role of *Cacna1b* during early postnatal development remains unclear, in adult mice, systemic *Cacna1b*-deficiency results in a hyperactive phenotype and alterations in vigilance state transitions(21, 30). Recent transcriptomic and chromatin accessibility studies in the preoptic area of the hypothalamus have revealed that sensory input shapes the postnatal maturation of neuronal subtypes, including the refinement of receptor expression patterns critical for cell-cell communication(31). Thus, early-life physical contact with the mother likely plays a critical role in shaping synaptic transmission in the hypothalamus, in part through the regulation of specific calcium channel subtypes. This process may establish experience-dependent pathways through which affiliative stroking stimulation can promote sleep.

The present study investigated the short-term effect of maternal deprivation during infancy and demonstrated that the calming effect induced by back stroking is a developmentally transient response. However, maternal deprivation may show a long-term effect in social behavior, anxiety and depression-like behavior and innate behaviors, such as parental behaviors. Given that repeated back stroking-induced sleep may facilitate the maturation of sleep-wake patterns, it is possible that artificially reared mice may exhibit sleep and circadian abnormalities in adulthood. In humans, early-life experiences involving comforting physical contact with caregivers are known to shape both psychological well-being and physical health in later life. Individuals who have had limited exposure to positive and affectionate touch during early development, such as those who experienced neglect or abuse, often exhibit reduced sensitivity to the social and emotional aspects of affiliative tactile interaction in adulthood(32). Although skin-to-skin contact and affiliative tactile stimulation between parents and infants are already practiced in clinical settings(33), our findings provide a neurobiological basis for these interventions. Parental touch may help maturation of hypothalamic neural systems and promote systemic calming, potentially supporting the development of stress resilience. Future understanding how peripheral tactile stimulation contributes to systemic calming through neural mechanisms may advance our understanding of pediatric conditions such as sensory hypersensitivity, hyposensitivity, and anxiety, and may inform the development of novel therapeutic approaches.

## Materials and Methods

### Human participants

Mother-infant pairs were recruited through advertisements distributed at regular events for postpartum parents at Toho University Omori Medical Center and at local childcare support facilities. All participants were Japanese. None of the participants had any serious physical or mental illnesses. All mothers were either full-time homemakers or on maternity leave at the time of the study. The mother’s mean age was 34 ± 4.34 years. Infants who cried before or during the experiments were excluded from the study. A total of 15 infants participated in each of the back and abdomen stroking tasks. Of the 15 infants initially recruited, two boys began to cry during the back-of-head stroking task, which was conducted after the abdomen and back stroking, and were therefore excluded.

This resulted in a final sample of 13 infants for that task. The age and sex distribution of infants in the abdomen and back stroking tasks was 1.30 ± 1.43 years (7 males, 8 females). For the back-of-head stroking task, data from 13 infants (1.20 ± 1.15 years; 5 males, 8 females) were included in the final analysis. All procedures were approved by the Ethics Committee of the Faculty of Medicine at Toho University (Approval Protocol ID# A24085).

### Stroking experimental procedure in human infants

The studies were performed between 10:00 A.M. and 2:00 P.M. in the rectangular room (263 cm x 377 cm) at approximately 25 °C as described in our previous study(34). When mothers and infants arrived at the university laboratory, the experimental procedure was explained to mothers, and they signed an informed consent form. We also obtained consent to use privacy-protected photos and videos by hiding participants’ faces. The mothers did not eat, drink or feed their infants 30 minutes before the experiment. The mothers did not use any fragrance and a wristwatch. Before the experiment, the mothers changed clothes, into plain shirts with short sleeves provided by the experimenters. To enhance the transmission of tactile stimulation from the mothers, the infants participated in this experiment wearing only diapers and baby underwear. Initially, to minimize stress for the infants, they can freely choose between sitting facing their mother on her lap or having both the mother and infant sitting facing forward (Fig. 1). If the infant starts by sitting facing the mother, the task begins with the mother stroking the infant’s back or the back of the head for 1 minute. This is followed by stroking the back of the infant’s head or back for another 1 minute, and finally, the infant is repositioned to sit facing forward, where the mother strokes the abdomen for 1 minute to complete the measurements. On the other hand, if the infant starts by sitting facing forward, the task begins with stroking the abdomen for 1 minute, followed by repositioning the infant to sit facing the mother, where the task of stroking the back or the back of the head for 1 minute is carried out. Before and after each of the three tasks of stroking different body parts, a 1-minute interval was inserted where the mother and infant sat without any stroking. During the measurement, the mother did not rock the infant, make eye contact, or talk to them.

### Behavioral and heart rate measurements in human infants

Two observers (S.Y. and M.Y.) independently recorded infants’ behaviors, including vocalizations and movements of the head, upper body, and lower body during each task. Using a stopwatch, they measured the duration of movement for each body part and calculated its proportion relative to the 1-minute task duration. Inter-observer agreement (S.Y. and M.Y.) was assessed using Cohen’s kappa, showing statistically acceptable reliability (k = 0.84 for head movement, k = 0.92 for upper body movement, and k = 0.85 for lower body movement).Three disposable ECG electrode patches (Nihon Kohden, Japan) were attached to the infants’ chests, following the method described in our previous study(34). ECG signals were continuously recorded throughout the experiment. All sessions were videotaped from the front using Handycam camcorders (Sony, Japan) and BIMUTAS-Video software (KISSEI COMTEC, Japan). ECG signals were sampled at 1000 Hz (Nihon Kohden). Time-domain parameters, including mean R-R interval (RRI) and mean instantaneous heart rate, were calculated using R version 4.4.2. These parameters were compared between the baseline period (1 minute prior) and during each stroking task (back, back of the head, and abdomen).

### Mice

All animal procedures were conducted in accordance with the Guidelines for Animal Experiments of Toho University and were approved by the Institutional Animal Care and Use Committee of Toho University (Approval Protocol ID #24-557). Breeding pairs of C57BL/6J (B6) or ICR mice were obtained from Japan SLC and CLEA Japan. Mice were raised in our breeding colony under controlled conditions (12 h light/dark cycle, lights on at 8:00 A.M., 23 ± 2°C, 55 ± 5% humidity, and ad libitum access to water and food). Mouse delivers were monitored twice daily, at approximately 10:00 and 18:00. The day on which newborn pups were first observed was designated as the day of birth (postnatal day (PND) 0). Within PND 1 to 2, litters were culled to six pups for B6 mice and to 14 pups for ICR mice. All experiments were performed from 8:30 to 12:30. In the present study, no significant differences were observed in physiological or behavioral measurements between male and female pups. Therefore, data from both sexes were combined for analysis.

### Assessment of maternal licking behavior

On PND 2, B6 pups were gently removed from the home cage, and three marks were placed on their back and one on their chest using a special marker (ZEBRA, Japan). The pups were then returned to the home cage with their dam. The marker used could not be removed by rubbing with dry cloth or paper but could be erased with a moist cloth or paper, allowing for evaluation of the extent of maternal licking. Approximately four hours later, the pups were again removed from the home cage, and the visibility of each mark was photographed alongside a ruler for quantification. To evaluate maternal licking patterns on the pup’s body, images were converted to grayscale, and mean brightness values within defined regions of interest were compared between pups immediately after marking and those after a 4-hour period in the home cage. Brightness values were quantified using ImageJ software (version 1.50i, NIH).

### Preparation of narrow enclosure and stroking procedure

A soft packing foam sheet was placed on top of a heating pad, and both ends were clipped to create a narrow enclosure approximately 6.5 cm in height and 8.5 cm in width. A new foam-sheet enclosure was prepared for each individual mouse pup and not reused. Since mouse pups typically remain in close physical contact with their littermates—a behavior known as huddling, soft pressure applied to both sides of the body may mimic this huddling behavior, helping pups stay within confined spaces(35). B6 and ICR pups were placed inside this setup at PND14 and PND13, respectively. After a 1-minute resting period, the experimenter gently stroked the pup’s pinnae for 1 minute and then left the pup undisturbed in the narrow enclosure for 3 minutes (no-stroking condition). The pup’s pinnae were then stroked again for 1minute, followed by 3 minutes of continuous back stroking with a soft paintbrush (stroking condition). Since mouse dams are known to thoroughly lick their pups to remove milk residues and feces that are firmly adhered to the pups’ bodies, maternal licking likely transmits tactile pressure not only to the surface of the skin but also to deeper subcutaneous tissues. To emulate this, the experimenter applied brush strokes with sufficient pressure to visibly move the pup’s back skin.

### EEG/ECG electrode implantation surgery

Male and female pups were implanted with EEG/ECG electrodes under anesthesia with isoflurane at PND13 in B6 mice and at PND12 in ICR strain. All surgical procedures were performed between 17:30 and 18:30. Prior to surgery, pups were fed soft milk agar using a spatula (15 g milk powder in 0.4% agarose; MARUKAN, Japan). The same milk agar was then placed in the postoperative cage. This procedure prevented deterioration in physical condition by the following morning, allowing for successful EEG/EMG recordings. After surgery, the pups were transferred to a new cage without the dam, equipped with a heating pad and a soft milk-agar block for weaning support. EEG and ECG recordings were conducted the following morning between 08:30 and 11:30. To avoid maternal rejection or dislodging of the head-mounted electrodes—caused by excessive licking or neglect in response to the protruding electrode pins—the pups were not returned to the home cage with the dam. The EEG/ECG electrode consisted of a four-pin gold-plated metal header with a 1.27 mm pitch. Two of the pins were connected to miniature screw electrodes (screw thread diameter: 1 mm; shaft length: 0.5 mm) via flexible Teflon-coated stainless-steel wires and were used for EEG recordings. The remaining two pins were connected to flexible Teflon-coated stainless-steel wires (1 cm and 7 cm in length, respectively) for ECG recordings. EEG electrodes were implanted epidurally over the parietal area (0.9 mm anterior to lambda, 0.9 mm lateral to the midline) referenced to the cerebellum area. For ECG recordings, one wire was inserted into the temporal muscle, and the other was tunneled subcutaneously and wound around the intercostal muscles near the heart. Both the EEG and ECG electrodes were affixed using a high-performance instant adhesive.

### EEG/ECG recording and analysis

On the morning following surgery, each pup was placed in a narrow enclosure, and EEG and ECG signals were recorded at a sampling rate of 500 Hz using the VitalRecorder system (Kissei Comtec, Matsumoto, Japan). The onset and offset of both the no-stroking and stroking periods were annotated based on EEG and ECG signals, as well as synchronized video data displayed via SleepSign software (Kissei Comtec, Matsumoto, Japan). Epochs showing slow-wave activity accompanied by behavioral quiescence were extracted as sleep-like states. EEG and ECG signals were exported from SleepSign in CSV format. The EEG data were subsequently imported into Spike2 software (Cambridge Electronic Design). Electromyography components were extracted using a 130–250 Hz bandpass filter (extEMG). We confirmed that the 130–250 Hz extEEG signal in mouse pups was nearly identical to that of EMG (Fig. S1). Delta components were extracted using a 1–4 Hz bandpass filter. Both extEMG and EEG signals were rectified and visualized for annotating the onset and offset of each event. Epochs previously identified as sleep-like in SleepSign corresponded to NREM sleep time windows in Spike2, during which the extEMG signal was nearly absent and delta power was elevated.

ECG data were processed using Microsoft Excel and R (version 4.4.2) software to calculate R-R intervals (RRI), and average heart rates were determined for the no-stroking, stroking, and NREM periods.

### Measurement of plasma corticosterone and oxytocin

To examine plasma corticosterone and oxytocin levels, three experimental groups were established: (1) pups isolated for 30 minutes inside the foam sheet enclosure, (2) pups continuously stroked on the back during the 30-minute isolation period in the enclosure, and (3) pups that remained in the home cage with their dams without disturbance. At the end of the experimental period, blood samples were collected by decapitation. Blood was collected into 1.5 ml microtubes containing 5 μl of heparin (1 unit/ μl). The samples were centrifuged at 2000 × g for 10 minutes at 4 °C, and the resulting plasma was transferred to new tubes. Plasma corticosterone and oxytocin levels were quantified using AssayMax Corticosterone ELISA kit EC3001-1 (ASSAYPRO, MO, USA) and Oxytocin Enzyme Immunoassay Kit K048 (ARBOR ASSAYS, MI, USA), respectively.

### Preparation of brain sections for histological analyses

Mouse pups were anesthetized with isoflurane and then transcardially perfused with 4% paraformaldehyde (PFA) in phosphate-buffered saline (PBS) for 2-4 h after the start of the light phase. The brains were postfixed in 4% PFA at 4°C overnight, followed by cryoprotection in 30% sucrose in PBS for 2 days, embedded in O.C.T. Compound (Sakura Finetek, Tokyo, Japan), and stored at −80°C. The frozen brains except olfactory bulbs were cryosectioned coronally at a thickness of 50 μm. The sections were stored in an antifreeze solution (0.05 M phosphate buffer, 30% glycerol, 30% ethylene glycol) at −25°C until use. To confirm the reproducibility of the staining, sections from two or three pups from different litters were used for each staining combination.

### *In situ* hybridization chain reaction using short hairpin DNAs and immunohistochemistry

To detect *c-Fos, Vglut2,* and *Vgat* mRNA expression, gene-specific DNA hairpins were prepared, and in situ hybridization chain reaction (isHCR) was performed using short hairpins, as described in previous studies (36). For *c-Fos* probes, 18 target sequences were selected from the full-length cDNA using a homology search by NCBI BLASTn (https://blast.ncbi.nlm.nih.gov) (Table S6). For combined isHCR and immunohistochemistry (IHC), tissue sections were blocked with 0.8% Block Ace (Dainihon Seiyaku) in PBST, followed by overnight incubation at 4 °C with anti-Cav2.2 antibody (1:600; ACC-002, Alomone Labs) diluted in 0.4% Block Ace in PBST. After three washes with PBST, the sections were incubated for 1 h at room temperature with goat anti-rabbit IgG conjugated to Alexa Fluor 568 (ab175471) and Hoechst 33342 (1 µg/ml). Sections were then mounted on glass slides and coverslipped using antifade mounting medium (VECTASHIELD Vibrance). For brightfield IHC, we employed a biotin-conjugated anti-rabbit secondary antibody (1:1,000), an avidin-biotin-peroxidase complex (ABC kit; Vector Laboratories, CA, USA), and DAB substrate solution (Nacalai Tesque, Kyoto, Japan), following our previously described protocol(37). Brightness values were quantified using ImageJ software (version 1.50i, NIH).

### Artificially reared pup

Artificially reared groups were established using ICR strain, following previously published protocols(38). To mitigate litter-specific effects, experimental data were collected using artificially reared and maternally reared pups derived from at least three different litters. In both the artificially reared and the maternally reared groups, litters were culled to 14 pups per litter on PND1. Pups in the artificial rearing group were housed in small plastic containers maintained at high temperature and humidity, with 6–7 pups per container. Each pup was marked on the back with a pen for identification. From 08:30 to 24:30, pups were fed every 4 hours. Prior to each feeding, gentle stimulation of the anogenital region was performed to induce urination and defecation, followed by weighing. After feeding, the pups were weighed again and returned to the container. Although milk intake varied among individuals, feeding was terminated when pups voluntarily stopped suckling on the artificial nipple, indicating satiety. During feeding, contact between the experimenter (wearing gloves) and the pups was kept to a minimum; no stroking or fur grooming was performed. On PND13, 30 minutes after the first morning feeding, EEG recordings and plasma corticosterone CORT measurements were conducted.

### RNA-Seq

At PND13, AR and MR pups were perfused with 20% RNAlater (ThermoFisher, MA, USA) diluted in ice-cold PBS, after which the brains were removed. Coronal brain slices of 1-mm thickness were prepared using a brain slicer (Muromachi, Tokyo, Japan), immersed in fresh RNAlater, and stored at −80°C until further processing. Slices four through seven contained the hypothalamic region, including the medial preoptic area and the retromammillary nucleus. Under a stereomicroscope, the hypothalamic area surrounding the third ventricle was dissected using a surgical scalpel along a square mold (approximately 2 mm × 3 mm) created with a bent 33G needle. Total RNA was extracted using the NucleoSpin RNA kit (Takara Bio, Shiga, Japan) from four MR and three AR pups, each derived from three different litters.

RNA quality was verified via spectrophotometric analysis, and next-generation RNA sequencing (RNA-Seq) was performed (Takara Bio). All R scripts used for preprocessing, normalization, dimensionality reduction, and differential expression analysis are publicly available at https://github.com/Makoto-Kashima/mouse20250709/blob/main/seurat_TCC_pipeline.R. The code includes pipelines for performing Seurat (version 5.3.0)(39)-based normalization and visualization, and conducting differentially expressed gene analysis using the TCC package (version 1.48.0)(40).

### Quantitative RT-PCR

Total RNA, as described above, was used for cDNA synthesis using the SuperScript III First-Strand Synthesis System (Invitrogen, MA, USA). Real-time quantitative PCR was performed using SYBR Green qPCR Master Mix (Thermo Fisher Scientific) following the manufacturer’s protocol. The following primers were used: *Gapdh* forward, 5′-AACTTTGTCAAGCTCATTTCCTGGT-3′; *Gapdh* reverse, 5′-GGTTTCTTACTCCTTGGAGGCCATG-3′; *Cacna1b* forward, 5′-CTCACGTCTCTTGTGGTCTTG-3′; *Cacna1b* reverse, 5′-TCCAGATTTGGTCCCTGTTATG-3′. All reactions were run in duplicate on an Applied Biosystems 7500 Fast Real-Time PCR System. Relative gene expression was calculated using the comparative Ct method (ΔΔCt method). The expression level of *Cacna1b* gene was normalized to that of *Gapdh* gene, and fold changes were determined using the formula 2^–ΔΔCt.

### AAV preparation

Three distinct target sequences for *Cacna1b*, as recommended by the Broad Institute (Table S2), were inserted into the pAAV-hSyn1EmGFP-3x-miR-shRNA*(Cacna1b*)-WPRE-bGHpA vector to generate a short hairpin RNA (shRNA) plasmid construct (sh*Cacna1b*), which was subsequently packaged into adeno-associated virus 8 (AAV8) particles (VectorBuilder). The titer administrated to mice in this study was 1.31 x 10^13^ GC/ml. A nonsilencing scrambled sequence was used to generate a control shRNA vector (shCtrl) (Table S7).

### Stereotaxic surgery

The stereotaxic apparatus was adapted from Chen et al(41). On PND1, pups were placed on crushed ice for 3–4 minutes. Complete cryoanesthesia was confirmed by cessation of all movements and a change in skin color from pink to purple. Cryoanesthetized pups received bilateral viral injections using glass capillaries connected to a Nanoject II system (Drummond, PA, USA). Prior to injection, the skin surface on the pup’s head was slightly pricked with a 33-gauge needle. Approximately 300 nL of virus (50.6 nL per injection, administered six times) was slowly injected into each hemisphere (coordinates relative to the confluence of sinuses as lambda: anteroposterior +0.5 mm, mediolateral ±0.5 mm, dorsoventral −3 mm). The capillary was withdrawn slowly after waiting for 1 minute post-injection. Pups were then placed on a warming blanket until fully recovered. The pups were tattooed on a paw or foot using a tattooing device (Natsume, Tokyo, Japan) for individual identification and then returned to their home cage. All injections were conducted between 09:00 and 12:00.

### Statistics

The data were preprocessed and visualized with Microsoft Excel and R version 4.4.2. All statistical analyses were conducted using R. We performed One-way ANOVA, paired t-

test, repeated-measures ANOVA, Welch’s t-test. Significance was set at p < 0.05 after p-value correction by Holm’s method.

## Data availability

The RNA-seq data have been deposited in the Gene Expression Omnibus (GEO) under accession number GSE305688. All other data supporting this study are available from the corresponding authors without restriction.

## Acknowledgments

We appreciate H.F. lab members, Dr. Kunio Sugiyama in Omori hospital at Toho university. JSPS KAKENHI (23K06798).

## Author contributions

S.Y. and H.F. designed the research and wrote the paper; S.Y. performed the research; S.Y., A.H., T.M., Y.T., K.N., K.M., M.W., and Y.H. contributed to establishing the experimental setup and/or analytical tools; S.Y. and M.K. analyzed the data. S.Y. and H.F. acquired funding.

