## Supplemental Figures and Tables for "Maternal care during early development is necessary for the acquisition of a calming response to back stroking"

This PDF file includes:
Figures S1 to S4
Tables S1 to S7

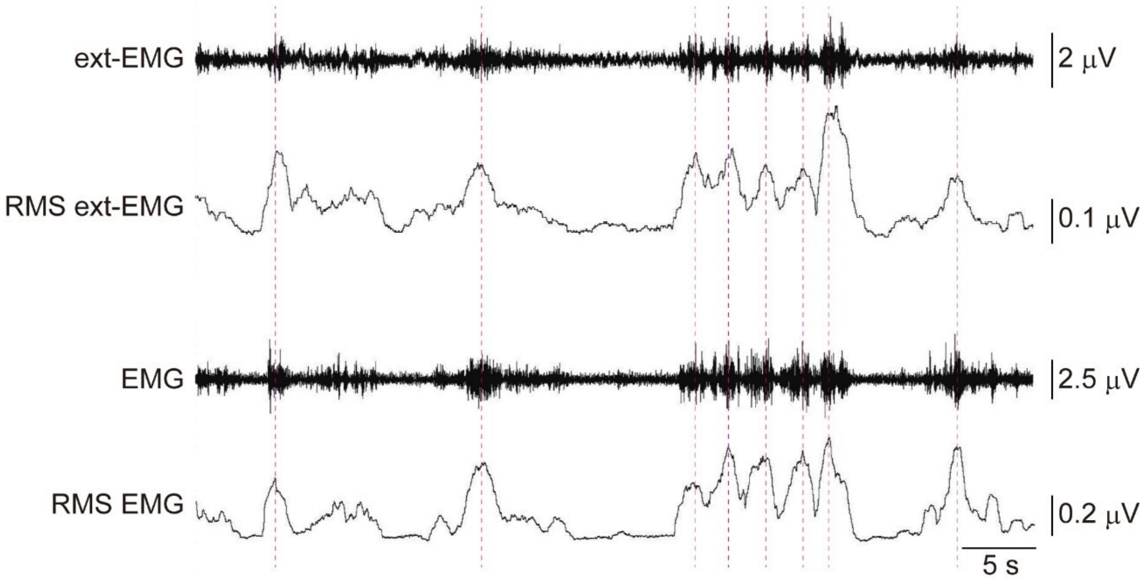


**Figure S1. Comparison of EMG extracted from EEG and EMG**

The top two panels show the EEG signal filtered in the 130–250 Hz range (ext-EMG, top) and its rectified RMS signal (RMS ext-EMG, second from the top). The bottom two panels show the raw EMG signal recorded from posterior neck muscles (EMG, second from the bottom) and its rectified RMS signal (RMS EMG, bottom).

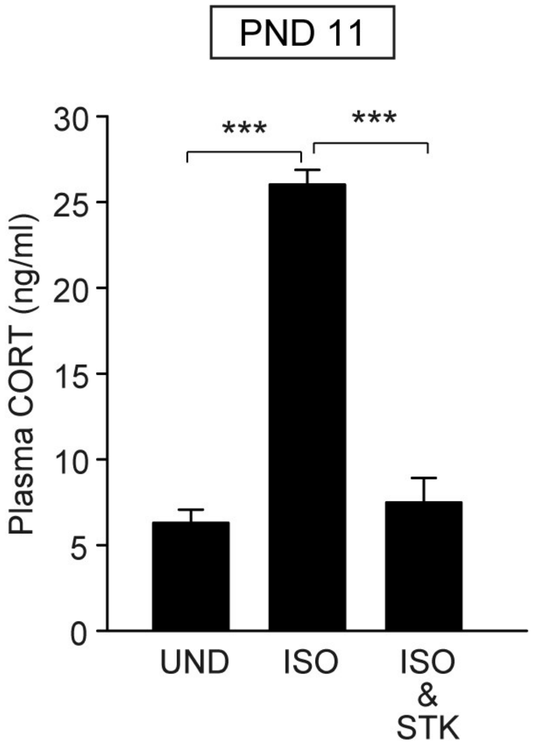


**Figure S2. Plasma CORT level in PND11 pups.**

Comparison of plasma CORT levels among undisturbed, isolated, and stroked during isolation conditions at PND11. n = 3 in each group. ***: $p < 0.001$

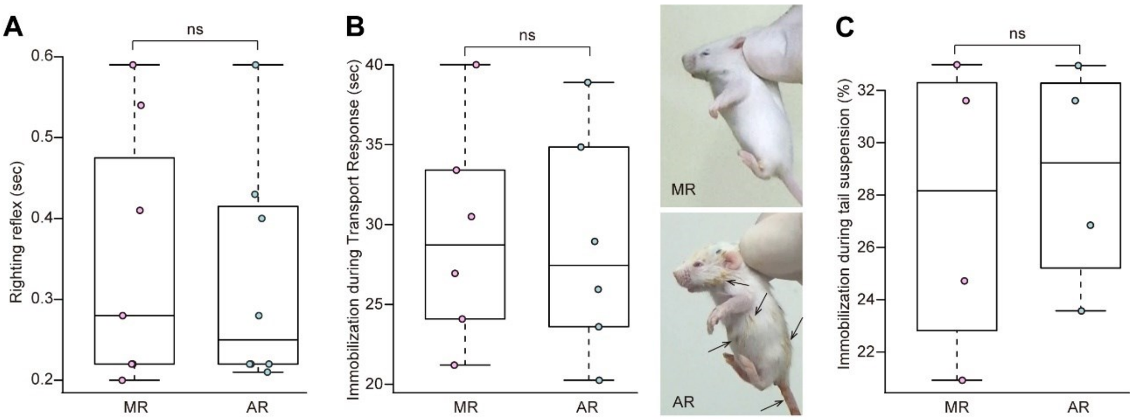


**Figure S3. Comparison of whole-body movements between AR and MR mice.**

(A) The duration of righting reflex in MR and AR pups at PND12. (B) Immobilization time during Transport Response in MR and AR pups at PND13 (left) and their general appearance (right). Arrows indicate fecal or milk residues adhered to the fur surface. (C) Immobilization time during 6-min tail suspension test in MR and AR mice at 8 weeks. (A, B) n = 6 (m:3, f:3). (C) n = 4 (m:2, f:2).

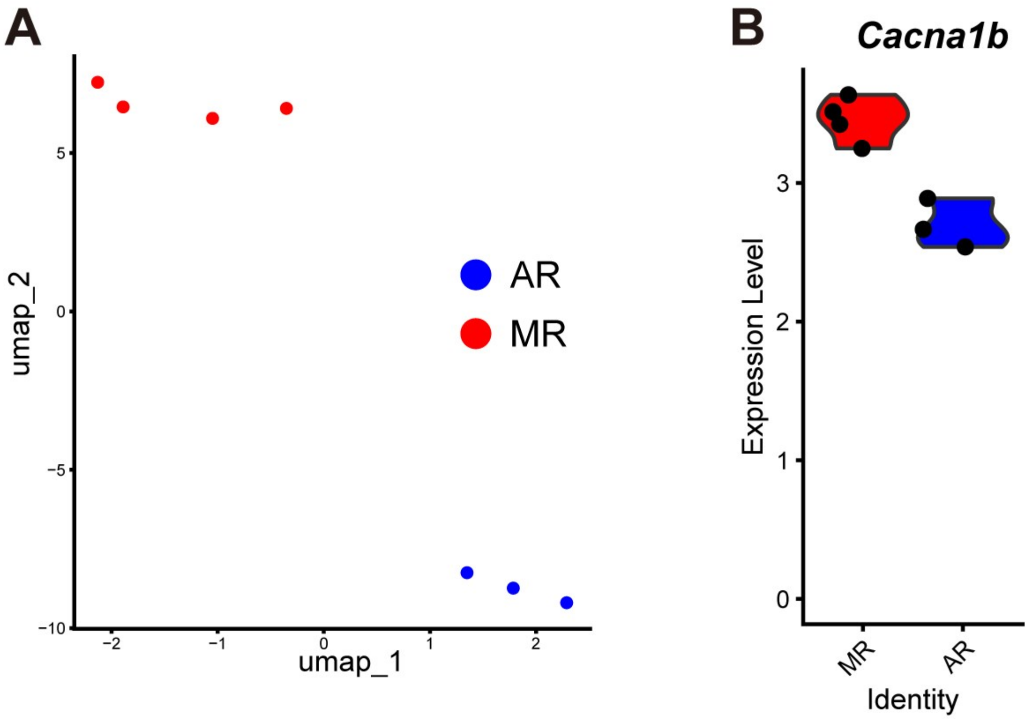


**Figure S4. UMAP-based clustering reveals transcriptomic differences between MR and AR pups**

(A) UMAP visualization of RNA-seq data reveals that MR and AR hypothalamic samples form distinct clusters, suggesting group-specific transcriptomic signatures. Each dot represents a sample. (B) The expression level of Cacna1b was lower in AR pups than in MR pups.

| Stroking area | Measurement index | t | df | p |
|---|---|---|---|---|
| Back of the head | Head movement | -0.60 | 6.88 | 0.57 |
| | Upper body movement | -0.040 | 5.95 | 0.97 |
| | Lower body movement | 0.35 | 4.79 | 0.74 |
| | Hear rate | 0.38 | 9.76 | 0.71 |
| Lower abdomen | Head movement | 1.19 | 9.14 | 0.26 |
| | Upper body movement | -0.68 | 9.38 | 0.51 |
| | Lower body movement | -0.14 | 12.95 | 0.87 |
| | Hear rate | -1.24 | 12.72 | 0.24 |
| Middle back | Head movement | -0.45 | 10.53 | 0.66 |
| | Upper body movement | 1.00 | 12.19 | 0.34 |
| | Lower body movement | 0.58 | 13 | 0.57 |
| | Hear rate | -0.41 | 12.77 | 0.69 |

**Table S1.** Results of t-tests comparing changes between males and females in each stroked body area.

| Mouse type (Related Figure) | Measurement index | t | df | p |
|---|---|---|---|---|
| MR B6 (Fig. 3C) | RMS EMG | -0.81 | 3.18 | 0.47 |
| | Heart rate | -0.66 | 3.79 | 0.54 |
| | RMS Delta power | 0.23 | 3.62 | 0.83 |
| MR ICR (Fig.5C) | RMS EMG | -0.72 | 1.39 | 0.57 |
| | Heart rate | -2.43 | 2.80 | 0.099 |
| | RMS Delta power | 1.98 | 2.016 | 0.18 |
| AR ICR (Fig.5D) | RMS EMG | 0.034 | 2.17 | 0.98 |
| | Heart rate | -1.67 | 3.66 | 0.18 |
| | RMS Delta power | -0.92 | 2.24 | 0.44 |
| ShCtrl (Fig.6H) | RMS EMG | -2.19 | 2.02 | 0.16 |
| | Heart rate | -3.04 | 1.13 | 0.18 |
| | RMS Delta power | 1.72 | 1.08 | 0.32 |
| ShCacna1b (Fig.6I) | RMS EMG | -0.28 | 2.18 | 0.81 |
| | Heart rate | 1.60 | 1.10 | 0.34 |
| | RMS Delta power | -1.46 | 3.33 | 0.23 |

**Table S2.** Results of t tests comparing changes in RMS EMG, heart rate, and EEG delta power between males and females in each experiment.

| Mouse type (Related Figure) | Sum of RMS Delta | t | df | p |
|---|---|---|---|---|
| MR B6 (Fig. 3D) | No-stroking | 0.15 | 3.96 | 0.89 |
| | Stroking | -0.68 | 2.89 | 0.54 |
| | NREM sleep | -0.35 | 3.86 | 0.75 |
| MR ICR (Fig.5E) | No-stroking | 2.48 | 2.59 | 0.10 |
| | Stroking | 0.85 | 1.64 | 0.50 |
| | NREM sleep | -1.21 | 2.96 | 0.31 |
| AR ICR (Fig.5F) | No-stroking | 0.62 | 2.95 | 0.58 |
| | Stroking | 0.30 | 1.44 | 0.80 |
| | NREM sleep | -1.52 | 2.22 | 0.26 |
| ShCtrl (Fig.6H) | No-stroking | -1.34 | 2.68 | 0.28 |
| | Stroking | -1.23 | 2.99 | 0.31 |
| | NREM sleep | -2.19 | 1.18 | 0.24 |
| ShCacna1b (Fig.6I) | No-stroking | 0.24 | 3.76 | 0.82 |
| | Stroking | -0.72 | 3.14 | 0.52 |
| | NREM sleep | 0.59 | 3.015 | 0.60 |

**Table S3.** Results of t-tests comparing sum of RMS delta between males and females.

| Mouse type (Related Figure) | Condition | t | df | p |
|---|---|---|---|---|
| MR B6 (Fig. 4B) | UND | -0.60 | 1.80 | 0.62 |
| | ISO | -0.34 | 1.76 | 0.77 |
| | ISO & STK | 1.99 | 2.00 | 0.19 |
| MR ICR (Fig.5G) | UND | 0.78 | 2.77 | 0.50 |
| | ISO | -0.52 | 1.39 | 0.67 |
| | ISO & STK | -1.19 | 1.16 | 0.42 |
| AR ICR (Fig.5H) | UND | 0.64 | 1.28 | 0.62 |
| | ISO | NA | NA | NA |
| | ISO & STK | -0.087 | 1.60 | 0.94 |

NA: No statistical test was performed for ISO AR pups (n = 3, m:1, f:2) due to limited sample size.

**Table S4.** Results of t-tests comparing plasma CORT between males and females.

| Gene Name | Gene Type | Transcript ID | Entrez Gene ID | Description |
|---|---|---|---|---|
| Gm15772 | processed pseudogene | ENSMUST00000118533.2 | - | predicted gene 15772 [Source:MGI Symbol;Acc:MGI:3805541] |
| Tuba1c | protein_coding | ENSMUST00000058914.10; ENSMUST00000230447.2 | 22146.0 | tubulin, alpha 1C [Source:MGI Symbol;Acc:MGI:1095409] |
| Rpl26 | protein_coding | ENSMUST00000073471.13; ENSMUST00000101014.9; ENSMUST00000128952.8; ENSMUST00000134403.2; ENSMUST00000138973.2; ENSMUST00000167436.3 | 19941.0 | ribosomal protein L26 [Source:MGI Symbol;Acc:MGI:106022] |
| Ahcy | protein_coding | ENSMUST00000054607.16; ENSMUST00000137242.2; ENSMUST00000146367.2 | 269378.0 | S-adenosylhomocysteine hydrolase [Source:MGI Symbol;Acc:MGI:87968] |
| Hspe1-rs1 | protein_coding | ENSMUST00000234633.2 | - | heat shock protein 1 (chaperonin 10), related sequence 1 [Source:MGI Symbol;Acc:MGI:1935159] |
| Gm49719 | lncRNA | ENSMUST00000231833.2 | - | predicted gene, 49719 [Source:MGI Symbol;Acc:MGI:6215192] |
| Hba-a1 | protein_coding | ENSMUST00000093209.4; ENSMUST00000142555.2 | 15122.0 | hemoglobin alpha, adult chain 1 [Source:MGI Symbol;Acc:MGI:96015] |

| | | | | |
|---|---|---|---|---|
| Folr1 | protein_coding | ENSMUST00000106981.8; ENSMUST00000106982.8; ENSMUST00000106983.8; ENSMUST00000106985.8; ENSMUST00000106986.9; ENSMUST00000123321.8; ENSMUST00000123630.8; ENSMUST00000124026.8; ENSMUST00000125298.2; ENSMUST00000126204.8; ENSMUST00000134145.8; ENSMUST00000140068.8; ENSMUST00000140584.2; ENSMUST00000150184.2; ENSMUST00000151706.8; ENSMUST00000155311.2 | 14275.0 | folate receptor 1 (adult) [Source:MGI Symbol;Acc:MGI:95568] |
| Hbb-bs | protein_coding | ENSMUST00000023934.8; ENSMUST00000131960.3; ENSMUST00000153218.2 | 100503605/15129 | hemoglobin, beta adult s chain [Source:MGI Symbol;Acc:MGI:5474852] |
| Cfap74 | protein_coding | ENSMUST00000050128.11; ENSMUST00000094408.10; ENSMUST00000105619.8; ENSMUST00000123952.9; | 544678.0 | cilia and flagella associated protein 74 [Source:MGI Symbol;Acc:MGI:1917130] |

| | | | | |
|---|---|---|---|---|
| | | ENSMUST00000129481.2; ENSMUST00000135407.8; ENSMUST00000144157.8; ENSMUST00000144625.2; ENSMUST00000151083.8; ENSMUST00000165947.3; ENSMUST00000238423.2; ENSMUST00000238620.2 | | |
| Rasd1 | protein_coding | ENSMUST00000062405.8 | 19416.0 | RAS, dexamethasone-induced 1 [Source:MGI Symbol;Acc:MGI:1270848] |
| Ubc | protein_coding | ENSMUST00000108707.3; ENSMUST00000136312.2; ENSMUST00000156249.2 | 22190.0 | ubiquitin C [Source:MGI Symbol;Acc:MGI:98889] |
| Gm4737 | protein_coding | ENSMUST00000059524.7 | 11615.0 | predicted gene 4737 [Source:MGI Symbol;Acc:MGI:3643647] |
| Ppp1r1b | protein_coding | ENSMUST00000078694.13; ENSMUST00000132443.8; ENSMUST00000133700.8; ENSMUST00000137634.2; ENSMUST00000147415.8; ENSMUST00000150762.8; ENSMUST00000152525.2 | 19049.0 | protein phosphatase 1, regulatory inhibitor subunit 1B [Source:MGI Symbol;Acc:MGI:94860] |

| | | | | |
|---|---|---|---|---|
| Sostdc1 | protein_coding | ENSMUST00000041407.7 | 66042.0 | sclerostin domain containing 1 [Source:MGI Symbol;Acc:MGI:1913292] |
| Matn2 | protein_coding | ENSMUST00000022947.7; ENSMUST00000163455.9; ENSMUST00000226766.2; ENSMUST00000227119.2; ENSMUST00000227759.2; ENSMUST00000228570.2 | 17181.0 | matrilin 2 [Source:MGI Symbol;Acc:MGI:109613] |
| Igsf1 | protein_coding | ENSMUST00000033442.14; ENSMUST00000072037.13; ENSMUST00000114891.2; ENSMUST00000114893.8; ENSMUST00000135492.2 | 209268.0 | immunoglobulin superfamily, member 1 [Source:MGI Symbol;Acc:MGI:2147913] |
| D3Ertd751e | protein_coding | ENSMUST00000026867.14; ENSMUST00000026868.13; ENSMUST00000108065.9; ENSMUST00000119572.8; ENSMUST00000120167.8; ENSMUST00000143841.8; ENSMUST00000146165.8; ENSMUST00000192193.6; ENSMUST00000192799.2; ENSMUST00000193075.6; | 73852.0 | DNA segment, Chr 3, ERATO Doi 751, expressed [Source:MGI Symbol;Acc:MGI:1289213] |

| | | | | |
|---|---|---|---|---|
| | | ENSMUST00000193228.6; ENSMUST00000194346.6; ENSMUST00000195030.2; ENSMUST00000195577.2; ENSMUST00000195882.6 | | |
| Cacna1b | protein_coding | ENSMUST00000041342.12; ENSMUST00000070864.14; ENSMUST00000100348.10; ENSMUST00000102939.9; ENSMUST00000114447.8; ENSMUST00000124183.2; ENSMUST00000125798.3; ENSMUST00000131861.2; ENSMUST00000133892.2; ENSMUST00000155356.4 | 12287.0 | calcium channel, voltage-dependent, N type, alpha 1B subunit [Source:MGI Symbol;Acc:MGI:88296] |
| Alas2 | protein_coding | ENSMUST00000066337.13; ENSMUST00000112715.2; ENSMUST00000134670.2; ENSMUST00000142474.2 | 11656.0 | aminolevulinic acid synthase 2, erythroid [Source:MGI Symbol;Acc:MGI:87990] |

**Table S5.** Description of 20 differentially expressed genes.

| Target | Sequence |
| --- | --- |
| *Cacna1b* | CCTTACTTTCGGGATCTTT |
| | GCGCATCATACAATGACAT |
| | GCCGTAATATCATGGGATT |
| Nonsilencing control | GCACTGGCGAGAGATGTAGTT |
| | GTGGTAATGTGCTTATTGTAT |
| | GCACAATACCGATAATCTGAT |

**Table S6.** Three distinct target sequences for sh*Cacna1b* and shCtrl.

| Probe name | First probe | Second probe |
| --- | --- | --- |
| cFos-1S10 | CGTCGGATGAAATGGTCGAAGTTTGGGGAAAGCCCG | TAGTCGGCGTTGAAACCCGAGAACAAAAGCCCATTAGAT |
| cFos-2S10 | CGTCGGATGAAGGGGAATGGTAGTAGGAAAGGCTGT | GAGCCCATGCTGGAGAAGGAGTCGGAAAGCCCATTAGAT |
| cFos-3S10 | CGTCGGATGAATGCGCAAAAGTCCTGTGTGTTGACA | AAAGTTGGCACTAGAGACGGACAGAAAAGCCCATTAGAT |
| cFos-4S10 | CGTCGGATGAAATGCTCTGCGCTCTGCCTCCTGACA | AGCTGCTCTACTTTGCCCCTTCTGCAAAGCCCATTAGAT |
| cFos-5S10 | CGTCGGATGAATCCGTTTCTCTTCCTCTTCAGGAGA | CCATCTTATTCCGTTCCCTTCGGATAAAGCCCATTAGAT |
| cFos-6S10 | CGTCGGATGAAGCAGACTTCTCATCTTCAAGTTGAT | AGCAGATTGGCAATCTCAGTCTGCAAAAGCCCATTAGAT |
| cFos-7S10 | CGTCGGATGAATCGGTGGGCTGCCAAAATAAACTCC | AAGGTCATCGGGGATCTTGCAGGCAAAAGCCCATTAGAT |
| cFos-8S10 | CGTCGGATGAAGCTTGGGCTCAGGGTCGTTGAGAAG | TGATGCTCTTGACTGGCTCCAAGGAAAAGCCCATTAGAT |
| cFos-9S10 | CGTCGGATGAAAAGGGTTCTGCCTTCAGCTCCACGT | GATGATGCCGGAAACAAGAAGTCATAAAGCCCATTAGAT |
| cFos-10S10 | CGTCGGATGAAATCTGGCACAGAGCGGGAGGTCTCT | TGCATAGAAGGAACCGGACAGGTCCAAAGCCCATTAGAT |
| cFos-11S10 | CGTCGGATGAACTCGGGCAGTGGCACGTCTGGATGC | GTGTTTCTCCTCTCTGTAATGCACCAAAGCCCATTAGAT |
| cFos-12S10 | CGTCGGATGAAAGGTCGACGGGAACCTTCGAGGGAA | GTTTCACGAACAGGTAAGGTCCTCCAAAGCCCATTAGAT |
| cFos-13S10 | CGTCGGATGAAGCAAGTCCTTGAGGCCCACAGCCTG | AGGACTGGAGGCCAGATGTGGATGCAAAGCCCATTAGAT |
| cFos-14S10 | CGTCGGATGAATCACTAGGAACAACACACTCCATGC | GCTCTACTAACTACCAGCTCTCAGGAAAGCCCATTAGAT |
| cFos-15S10 | CGTCGGATGAAGGTTAATTCCAATAATGAACCCAAC | AGCTGCACTAGATACAATCCAGCACAAAGCCCATTAGAT |
| cFos-16S10 | CGTCGGATGAACGCTATTGCCAGGAACACAGTAGGT | TAATATTGGTCGTTTCTAATTGGAAAAAGCCCATTAGAT |
| cFos-17S10 | CGTCGGATGAATGACGCTGAAGGACTACAGTACATG | CATGATCAGTAACATGACAATGAACAAAGCCCATTAGAT |
| cFos-18S10 | CGTCGGATGAAGAACATTCAGACCACCTCGACAATG | CGTTTTCATGGAAAACTGTTAATGTAAAGCCCATTAGAT |

**Table S7.** isHCR probe sequences for mouse *c-Fos* gene.
